# A hierarchical Bayesian approach to estimate endosymbiont infection rates

**DOI:** 10.1101/102020

**Authors:** Zachary H. Marion, Christopher A. Hamm

## Abstract

Endosymbionts may play an important role in the evolution of the Insecta. Bacteria such as *Wolbachia, Cardinium,* and *Rickettsia* are known to manipulate host reproduction to facilitate their own. Indeed, there are many documented cases where *Wolbachia* (Alphaproteobacteria: Rickettsiaceae) induces one of four manipulative phenotypes (cytoplasmic incompatibility, male killing, feminization, and parthenogenesis). The scale of infection among species has been a major subject of investigation, but quantification has been difficult because various approaches have yielded different estimates. One under-appreciated aspect of this problem arises when multiple—yet independent—samples are taken within a taxon. When independent samples within a taxon are treated as levels of a hierarchy, the problem is greatly simplified because data are partially pooled according to taxon. Here, we present a hierarchical Bayesian approach to estimate infection frequency where multiple independent samples were collected across several taxonomic levels. We apply this model to estimate the rates of infection for *Wolbachia* in the Lepidoptera, and then extend the model to account for phylogenetic non-independence. In addition, we highlight the current knowledge regarding *Wolbachia* and its effects within Lepidoptera. Our model estimates that the rate of endosymbiont infection for the Lepidoptera is approximately 12%. Given our limited knowledge regarding the phenotypes induced by these endosymbionts and the low infection rate, we urge caution when extrapolating the results of positive assays.

## 1 INTRODUCTION

Scientists have long known that bacterial endosymbionts inhabit insects. Many of these endosymbionts are maternally transmitted to offspring through the cytoplasm of the egg. *Wolbachia* (Alphaproteobacteria: Rickettsiaceae) was the first of these endosymbionts to be discovered when Hertig and Wolbach (1924) examined the adult ovaries and testes of *Culex pipiens* (hence the specific epithet of *Wolbachia pipientis*; Hertig, 1936). Some years later, Yen and Barr (1971) observed that male *C. pipiens* from one geographic area may not successfully reproduce with females from a different area and that reciprocal crosses could produce similar results; this phenomenon was given the name *cytoplasmic incompatibility*. A *Rickettsia-*like organism was determined to be the causative agent—*Wolbachia* (Yen and Barr, 1973). Recently, Hilgenboecker et al. (2008) estimated that about 20% of the Insecta are infected with *Wolbachia*, while Zug and Hammerstein (2012) placed that estimate at roughly 40%.

Today, researchers detect the presence of *Wolbachia* via the polymerase chain reaction (PCR), samples can be screened quickly and relatively inexpensively (Baldo et al., 2006; Simões et al., 2011). However, this development is relatively recent. Prior to the advent of PCR, *Wolbachia* infection was only confirmed through painstaking work that included electron microscopy and other microbiological techniques. Indeed, these methods were so laborious that they were only employed once a researcher had a prior reason (e.g., *male killing, feminization, parthenogenesis, cytoplasmic incompatibility*) to suspect the presence of the bacterium. We are aware of no cases in which exploratory assays for *Wolbachia* were conducted prior to the appearance of PCR.

Exploratory investigations for the presence of *Wolbachia* became feasible with the advent of PCR and Sanger sequencing. Yet few studies conducted the experimental work to determine if any reproductive manipulation was occurring. Careful experimental work is required to determine what (if any) phenotype is induced by an endosymbiont. The effects of *Wolbachia* infection are complex and depend on the interaction between the genomes of the endosymbiont and the host. For example, the phenotypic effects of one strain of *Wolbachia* may be very different if moved into another host (Rigaud et al., 2001; Hoffmann et al., 2011). Additionally, there may be extensive genomic differences between closely related strains (Ishmael et al., 2009). Although *Wolbachia* is most famous for being a ”reproductive parasite,” *Wolbachia* infections can often result in no manipulation at all (Hamm et al., 2014a; Zhang et al., 2010, 2013). Thus, infections do not necessarily cause reproductive manipulations.

The Lepidoptera (Arthropoda: Insecta) are among the best studied animal orders, containing ~ 160, 000 species in 124 families, representing approximately 13% of all described life (Regier et al., 2013; van Nieukerken et al., 2011). Because of historic interest in their physical beauty and their contemporary economic importance, the literature is replete with detailed information regarding their distribution and life history. In addition to basic scientific research, the Lepidoptera are also well represented on lists of endangered or threatened species (Hamm et al., 2014b). Yet certain groups of Lepidoptera have garnered the majority of attention, such as the butterflies (e.g. Nymphalidae, Lycaenidae and Pieridae) or groups of economically important pest species such as the Crambidae (which contains the Asiatic rice borer *Chilo suppressalis*) and Noctuidae (which contains the armyworms of the genus *Spodoptera*). This results in a bias towards certain groups and leaves most of the remaining families understudied.

Six species of Lepidoptera have been tested for the existence of a naturally occurring manipulative phenotype with evidence for *cytoplasmic incompatibility, male killing,* and *feminization* (Table 1). We note that the report of *male killing* in *Ephestia kuhniella* is a result of *Wolbachia* transfected from *Ostrinia scapulalis* (Fujii et al., 2001). Given the high level of interest in the Lepidoptera, understanding the role *Wolbachia* plays in the evolution of lepidopterans has received considerable attention. A vital first step towards this understanding is the estimation of *Wolbachia* infection rates across the order. Previous work on the estimation of *Wolbachia* infection frequency have employed maximum likelihood estimation. Ahmed et al. (2015) and Weinert et al. (2015) estimated *Wolbachia* infection frequencies for the Lepidoptera. These studies represent important steps in the estimation of *Wolbachia* infection frequency in the Lepidoptera, and our work here builds upon them. Our primary aim here was to develop a hierarchical partial-pooling strategy and account for phylogenetic non-independence at the family level.

**Table 1.** Published phenotypic effects of *Wolbachia* on Lepidoptera. Phenotype: MK = male killing, Fem = feminization, CI = cytoplasmic incompatibility. * = induced by transfection with *Wolbachia* strain from *O. scapulalis.*

| Species | Family | Phenotype | Reference |
| --- | --- | --- | --- |
| *Acraea encedana* | Nymphalidae | MK | Jiggins et al. (2000) |
| *Acraea encedon* | Nymphalidae | MK | Jiggins et al. (1998) |
| *Ephestia kuehniella* * | Pyrilidae | MK | Fujii et al. (2001) |
| *Eurema hecabe* | Pieridae | CI | Narita et al. (2007) |
| *Hypolimnas bolima* | Nymphalidae | MK | Dyson et al. (2002); Mitsuhashi et al. (2004) |
| *Ostrinia scapularis* | Crambidae | MK & Fem | Sugimoto and Ishikawa (2012) |
| *Ostrinia furnacalis* | Crambidae | Fem | Kageyama et al. (2002) |

Here, we develop a novel hierarchical Bayesian approach to estimate *Wolbachia* infection frequencies across the Lepidoptera. Our model explicitly accounts for issues that arise with real world data such as quantifying infection levels at different taxonomic levels. In a hierarchical Bayesian approach, a compromise via partial pooling occurs because lower levels of the hierarchy inform higher levels of the hierarchy, and vice versa. Therefore, When there is little information within a grouping (e.g., species with few observations), those estimates are pulled strongly (shrunk) towards the among-group mean. Conversely, parameter estimates for groups with high levels of information experience little shrinkage and instead inform the estimates for groups with less information. For example, there may be multiple observations of infection frequency collected from different populations within a species, often with disparate sample sizes. We do not consider it appropriate for these samples to be completely pooled, as that ignores population differences in infection frequency. Nor should observations within species be considered independent, because of shared ancestry. Similarly, there may be single samples collected from many different species within a family. In this case, individual sampling error should be accounted for when estimating family level infection rates. Finally, we consider that there has been a bias towards studying only a few families of the Lepidoptera. This uneven sampling can cause a few well-studied families to drive estimates of overall infection frequency. Each of these concerns can be specifically addressed using a hierarchical Bayesian model that incorporates phylogenetic relatedness among lepidopteran families.

## 2 MATERIALS & METHODS

### 2.1 Motivating data and previous analyses

Early studies on *Wolbachia* prevalence reported the frequency of infection for small groups of insects or arthropods (Jiggins et al., 2001; Werren and Windsor, 2000). More recent and sophisticated models of *Wolbachia* infection in the Lepidoptera used a likelihood-based approach to describe the distribution of *Wolbachia* infection across arthropods (Weinert et al., 2015) and the Lepidoptera specifically (Ahmed et al., 2015). Following Hilgenboecker et al. (2008), both studies used beta-binomial models to estimate the mean proportion of individuals infected within a given species. Both used the same distribution to calculate the incidence of infection as well, where incidence was the proportion of species infected above a threshold frequency (i.e., one infection in 1000 individuals, or 0.001; Weinert et al., 2015).

In the case of *Wolbachia*, tested insects may be either positive or not positive (a band of appropriate length when an electrophoresis gel is run, or no band, respectively). It is important to note that “not positive” is more appropriate than negative here because infections may have been missed for a number of reasons, including low density infections (Schneider et al., 2014). However, for the sake of simplicity, we will treat *Wolbachia* infection status as two mutually exclusive outcomes, (0 or 1; positive or not positive). This makes the question of infection a binomial sampling problem. The issue is the way that some models have accounted for uncertainty at each level. We will demonstrate this problem with two examples. First, let us assume that 200 individuals of a species are assayed for *Wolbachia*, and 100 of those tests are positive for infection. The mean estimate of infection is 0.5 and the 95% exact binomial confidence interval is 0.43–0.57. Next, consider two assayed individuals from a species where one tested positive. Here the proportion infected in this species is also 0.5, yet the 95% confidence interval is 0.01–0.99. It is clear that there is uncertainty around each estimate and that uncertainty varies with sample size. For this error to be properly incorporated into any estimate it must be treated at each level of the analysis (species, family, etc.), rather than pooled at the level of the entire study.

### 2.2 Data

We used the data compiled by Weinert et al. (2015), which contains records from thousands of individual sampling efforts across arthropods. Each row denotes one independent sampling event (though each row may contain data from multiple individual assays) and contained information on the arthropod family, genus, species, number of individuals assayed, number of individuals positive for infection, and endosymbiont genus. For this study, we only considered *Wolbachia* assays of Lepidoptera. All analyses were conducted in R (R Core Team, 2016, *v3.3*), and all data and code necessary to reproduce our results are freely available on Zenodo (http://doi.org/10.5281/zenodo.166803).

We used the Lepidoptera phylogeny of Regier et al. (2013) as a covariate to account for any influence of the relatedness among families in our analysis. This tree contained representatives from 115 of the 124 families in the order. The tree was pruned to remove duplicate entries at the family level and those not present in the Weinert et al. (2015) dataset. We made the tree ultrametric using the penalized likelihood method of Sanderson (2002) with tools from the *ape* package (Paradis et al., 2004). To incorporate phylogenetic history into the Bayesian model, we used the pruned ultrametric tree to create a series of phylogenetic correlation matrices. We constructed one matrix in which we assumed that *Wolbachia* infection status was distributed according to Brownian Motion (BM), a model of trait evolution that assumes closely related taxa share that trait due to common ancestry (Paradis, 2012). We also constructed matrices which assumed trait evolution followed an Ornstein-Uhlenbeck (OU) process, which places constraints around which a character evolves (Paradis, 2012). Relative to the BM, the OU model has two additional parameters: *θ* (the ”optimal” value for a character), and *α* (the rate at which *θ* moves towards *α*) (Paradis, 2012). The *α* value can range from 0 - 1; when *α* is 0 the model is effectively pure BM and becomes less so as *α* increases. We rescaled the phylogeny using three alpha values to examine their impact: *α* = 0.1 (similar to BM), *α* = 0.5, and *α* = 0.9 (very different than BM). Finally, we used an identity matrix (ones on the diagonal and zeros for the off-diagonal correlations) that assumed no phylogenetic correlation at all. We should note that all these correlation matrices (and the identity matrix) are accounting for relatedness at the family level only because of taxonomic incompleteness at the generic or species level. Because of this taxonomic incompleteness, we assume a star phylogeny for species within families.

### 2.3 Bayesian hierarchical models

For our hierarchical Bayesian model to estimate the probability of infection prevalence within and among members of Lepidoptera, each observation (*N* = 1037)—the number of *Wolbachia*-infected individuals—was nested within species (*S* = 411) and modeled as:

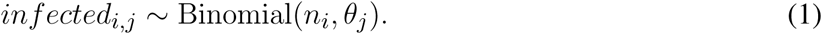

where *i* = 1, 2, …, 1037 and *j* = 1, 2, …, 419. Here *infected*_*i,j*_ indicates the number of infected individuals from the *i*th observation of the *j*th species, *n*_*i*_ is the total number of screened insects in observation *i*, and *θ*_*j*_ is the probability of infection for species *j*.

We then assumed the species-level probabilities of infection were normally-distributed with family-level means (*μ*_*k*_) and standard deviations (*σ*_*k*_) where *k* = 1, 2, …, 28 families. For computational efficiency, we used a non-centered parameterization of the normal (Papaspiliopoulos et al., 2007). The normal distribution is unconstrained, but is bounded between zero and one. Therefore the species-level *θ*s were logit transformed such that:

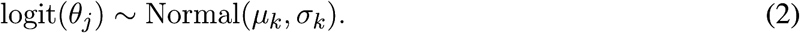

The mean (*μ*_*k*_) describes the average probability of infection within a family on the log-odds scale and can be back-transformed using the inverse-logit function.

The standard deviation (*σ*) measures how much variation in the probability of infection there is across species. If *σ* is small, then infection probabilities will be similar among species. Conversely, if *σ* is large, species-specific probabilities of infection will be more idiosyncratic. Data sparsity can be a problem in hierarchical models, especially for the estimation of scale parameters like variances. Because there were several species with few observations, we used shrinkage priors (Carvalho et al., 2009, 2010) for the species-specific *σ*s:

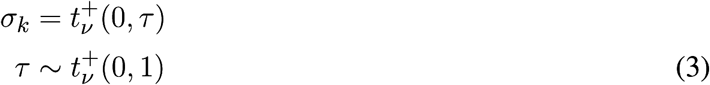

where 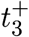 is half-Student-t distribution with *ν* = 3 degrees of freedom.

We modeled *μ*, the vector of log-odds infection probabilities for families using a multivariate normal distribution:

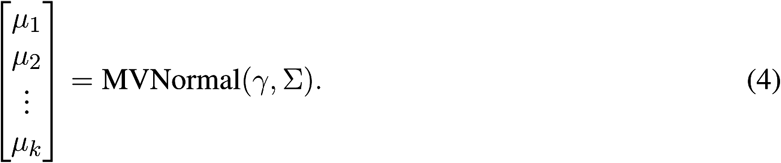

with the mean log-odds probability of infection across Lepidoptera (*γ*) and covariance matrix Σ. To account for phylogenetic non-independence among families, we constructed sigma as:

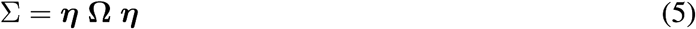

where ***η*** is a *k × k* diagonal matrix with the overall standard deviation on the diagonals and Ω is a *k × k* phylogenetic correlation matrix. We then put regularizing priors (to prevent overfitting) on both *γ* and *η*:

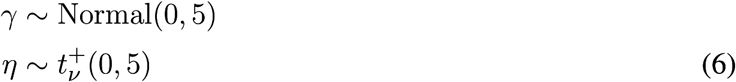

where, again, 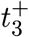 is half-Student-t distribution with *ν* = 3 degrees of freedom.

Posterior probabilities for model parameters were estimated using Markov chain Monte Carlo (MCMC) sampling in the Stan programming language (Carpenter et al., 2016) via the *RStan* interface (Stan Development Team, 2016). For each model, four MCMC chains were used with 5,000 iterations each. The first 2,500 iterations for each chain were adaptive and discarded as warm-up. We used several diagnostic tests to confirm that each model had reached a stationary distribution, including visual examination of MCMC chain history, calculation of effective sample size (ESS), and the Gelman-Rubin convergence diagnostic (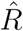; Gelman and Rubin, 1992; Brooks and Gelman, 1998). In particular, model convergence was assessed by inspecting the diagnostics of the log-posterior density. Model fit was also assessed by posterior predictive checks by simulating “new” data from the posterior distribution and plotting it against the original data (Gelman et al., 2013).

We used WAIC (the widely applicable or Watanabe-Akaike information criterion; Watanabe, 2010; Gelman et al., 2014) to compare models with different phylogenetic correlation matrices (e.g., Brownian motion vs. OU processes) using the *loo* package (Vehtari et al., 2016).

## 3 RESULTS

After filtering the Weinert et al. (2015) data to contain only Lepidoptera assayed for *Wolbachia* we retained 1037 independent samples from 411 unique species across 28 families, representing a total of 10860 individual assays (Figure 1a). Of these, 3607 samples from 163 species in 19 families were scored PCR positive for *Wolbachia*.

**Figure 1a.**
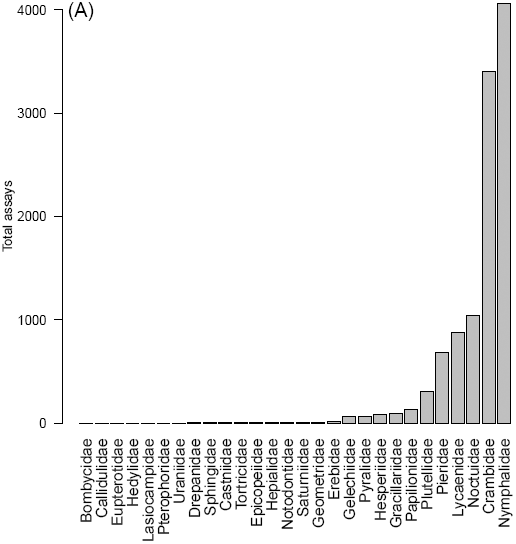
Total assays by family.

**Figure 1b.**
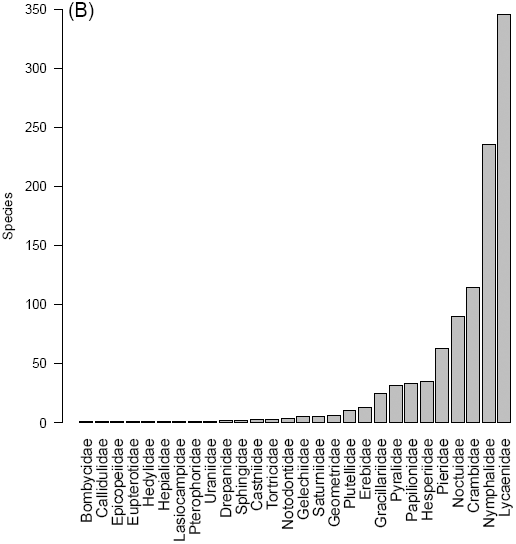
Number of species sampled by family

**Figure 1c.**
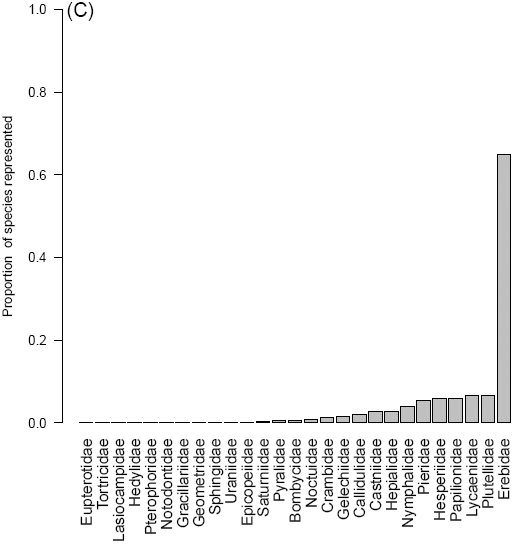
Proportion of sampling events per family

The 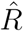 diagnostic for all parameters (including the log-posterior density) was 1.0, indicating that each model had reached a stationary posterior distribution. Visual assessment of the MCMC chain history confirmed this. Additionally, the effective sample size for the log-posterior density was *>* 2000 for all models. Predictive plots of the posterior means of the simulated “new” observations regressed against the original observations (Figure 2) resulted in tight concordance, suggesting the models were doing an excellent job at describing the data (all models: *R*^2^ = 0.91). If the model provides a perfect fit, the intercept of the slope should be zero and the slope should equal one. For all models, including the averaged consensus model (see below), the regression intercept was 0.41 (0.12 SE). Given that the data ranged from 0–255, the intercept is effectively zero. Additionally, the slope was 0.88 (0.008 SE), quite close to the theoretical optimum of one.

**Figure 2.**
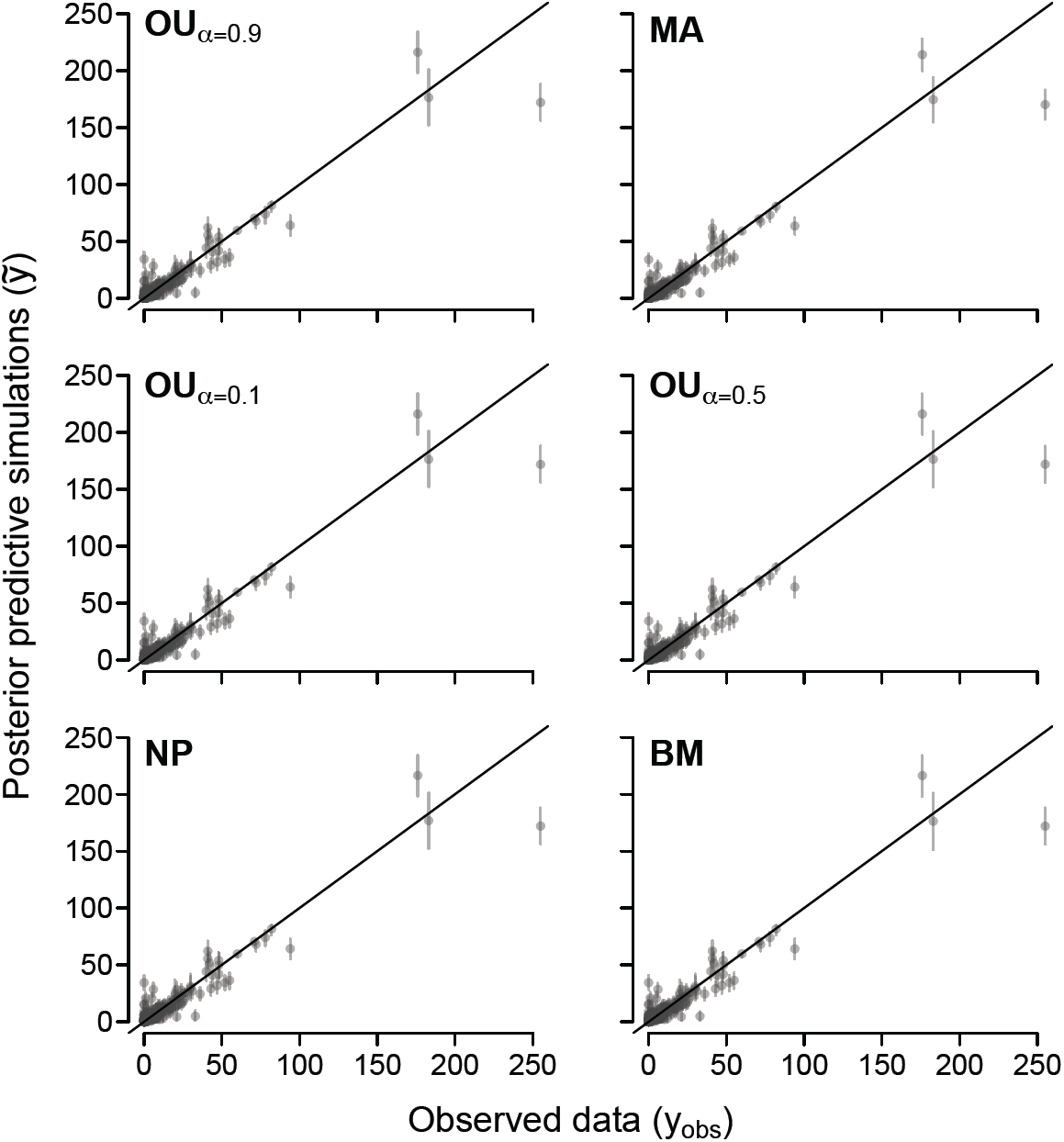
Plots of posterior predictive simulations (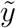) regressed against the observed data (*y*_*obs*_). Points are means of the posterior predictive simulations for each data point, while error bars around each point are the 95% Highest Density Intervals (HDI). Points are partially transparent to show where the majority of the data lie. Ideally, the observed data and the simulated data should have one-to-one correspondence and fall perfectly along a regression line with intercept of zero and slope of one (shown). Different panels represent models with different phylogenetic correlation matrices. NP = No Phylogenetic correction; BM = Brownian Motion; OU = Ornstein-Uhlenbeck with varying levels of *α* (0.1, 0.5, 0.9); MA = WAIC model weighted average.

All models, including those which contained phylogenetic correction, had similar WAIC scores with standard errors that completely overlapped (Table 2). We interpret this to mean that each model was in the same “family” of best models. Additionally, the parameter estimates were almost identical across models, and all models predicted a median infection frequency of ~ 12%. Rather than consider each model separately, we created a consensus model using model weighted averaging based on the Δ_*W AIC*_ scores (Table 2) to describe *Wolbachia* infection frequency in the Lepidoptera.

**Table 2.** Models with different phylogenetic correlation structures ranked according to WAIC and their respective model weights. The no-phylogeny model had an identity matrix (ones on the diagonal and zeros on the off-diagonals) in place of a correlation matrix. Smaller WAIC values indicate better estimates. Δ_waic_ is the difference between each WAIC and the lowest WAIC value. SE_waic_ and SE_Δ_ are the standard errors for WAIC and Δ_waic_ respectively.

| Model | WAIC | $\text{SE}_{\text{waic}}$ | $p_{\text{waic}}$ | $\Delta_{\text{waic}}$ | $\text{SE}_{\Delta}$ | weight |
| --- | --- | --- | --- | --- | --- | --- |
| OU: $\alpha = 0.1$ | 3,458.7 | 318.1 | 245.2 | 0.0 | | 0.79 |
| OU: $\alpha = 0.5$ | 3,462.8 | 319.7 | 245.7 | 4.1 | 2.55 | 0.10 |
| Brownian Motion | 3,464.1 | 320.3 | 246.6 | 5.4 | 3.44 | 0.05 |
| No Phylogeny | 3,464.9 | 320.2 | 247.9 | 6.3 | 3.07 | 0.03 |
| OU: $\alpha = 0.9$ | 3,465.9 | 320.1 | 247.1 | 7.2 | 3.13 | 0.02 |

Our estimate for the median *Wolbachia* infection frequency in the Lepidoptera was 12.1% (95% Highest Density Interval (HDI) = 0.045–0.33; Figure 3). Estimates of median family-level infection frequencies varied considerably with a positive association between sample size and HDI (Figure 4, Supplemental Figure 1). For example, the Nymphalidae (4060 specimens from 236 species) and Lycaenidae (878 specimens from 346 species) had relatively tight posterior distributions: median estimates of infection were 0.037 (95% HDI: 0.015, 0.082) and 0.24 (95% HDI: 0.13, 0.36) respectively. In, contrast families that had small sample sizes (e.g. Bombycidae, Hedylidae, and Lasiocampidae) generated larger intervals reflecting uncertainty in the estimates. For example, the HDI for Hedylidae was 0.06–0.8.

**Figure 3.**
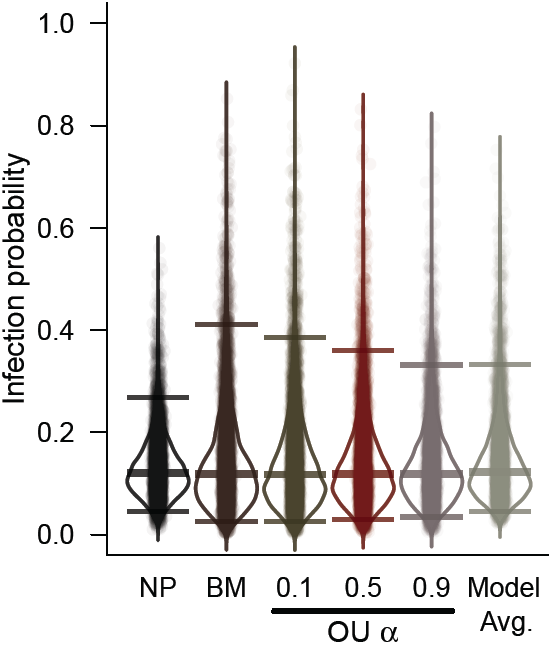
Posterior density plots for the average frequency of *Wolbachia* infection across Lepidoptera. Each posterior estimate is jittered and superimposed on the violins with transparency. Fatter regions of the violins indicate regions of higher posterior density, as do darker regions of jittered points. Horizontal bars indicate the median and upper and lower 95% Highest Density Interval (HDI). Models (L to R): NP = No Phylogenetic correction; BM = Brownian Motion; OU = Ornstein-Uhlenbeck with varying levels of *α* (0.1, 0.5, 0.9); Model Avg. = WAIC model weighted averaging.

**Figure 4.**
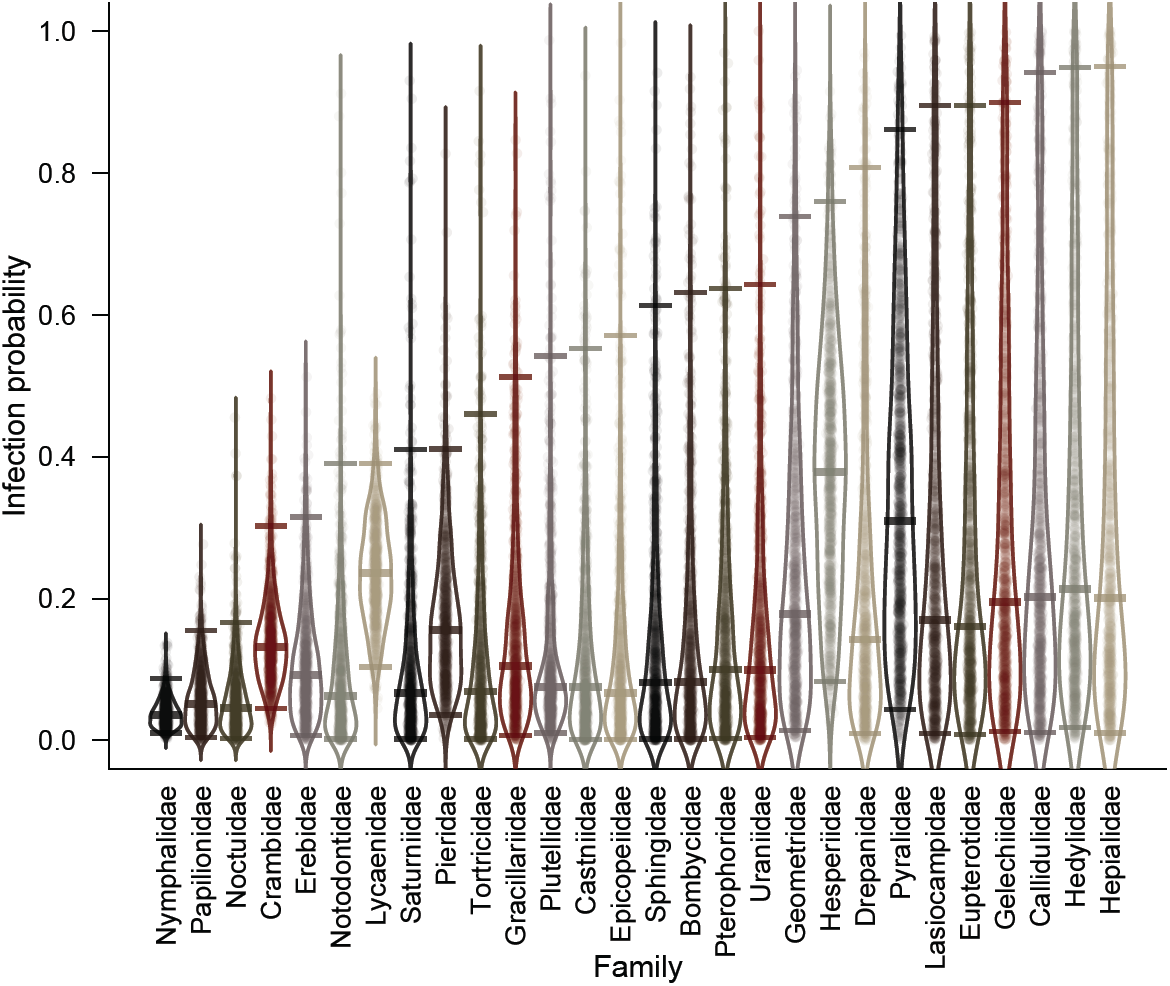
Posterior density plots for the average frequency of *Wolbachia* infection among 28 families of Lepidoptera. Each posterior estimate is jittered and superimposed on the violins with transparency. Fatter regions of the violins indicate regions of higher posterior density, as do darker regions of jittered points. Horizontal bars indicate the median and upper and lower 95% Highest Density Interval (HDI).

## 4 DISCUSSION

Our model predicts an average *Wolbachia* infection rate across lepidoptera of approximately 12%. As with Weinert et al. (2015), we consider that there are three main sources of bias in the dataset: 1) some species are represented by a single sample (Figure 1b), 2) there is a taxonomic bias in the data (Figure 1c), and 3) research may be focused on groups with known *Wolbachia* infections (e.g., Nice et al., 2009). Additionally, We consider a fourth source of bias: some families were extensively sampled among a small number of different species. This may bias the results towards a few members of an otherwise large family of Lepidoptera. We suggest that our estimates of infection rate are lower than previous research because our hierarchical Bayesian model accounts for these biases at each level and therefore may produce more reliable estimates.

We find it interesting that our median infection frequency estimates for the Lepidoptera do not significantly change when the model considers relatedness by incorporating phylogenetic information (Figure 3). Additionally, the model WAIC scores were within 8 units of one another and their standard errors completely overlapped. This implies that the models were all well within the same “family of best models” (Table 2). We interpret these results to indicate that our model is robust to the differential sampling present in the current data set. We consider our estimate for median *Wolbachia* infection rate for the Lepidoptera, and many of the family-level estimates, to be reliable given the limitations of the data. Estimates of infection frequency for those families with large sample sizes are likely accurate, but we must advise caution when interpreting some of the estimates for families with small sample sizes. In these cases where one sample has been assayed for an entire family (Bombycidae, Callidulidae, Eupterotidae, Hedylidae, Lasiocampidae, Pterophoridae, Uraniidae), the estimates presented in Figure 4 are strongly influenced by families with more complete sampling, and are shrunk towards the hyperprior—i.e., the overall probability of infection across Lepidoptera. Therefore, we have little faith in point estimates (i.e., mean, median) for these families and this is reflected in the large HDIs.

In addition to the lower estimate of *Wolbachia* infection for the Lepidoptera, our family level estimates were also lower than those reported by Ahmed et al. (2015) (we restrict our comparison here to those families with larger sample sizes). In these cases, as with our estimates of order level infection, the highest density intervals for our estimates were also much larger and we attribute this to the hierarchical manner in which error is handled. Using linear regression, we observed a strong association between the number of samples assayed per family and the range of the 95% HPD, where small sample sizes generated larger HPD ranges (*F*_1,26_ = 12.56, *P* = 0.0015).

Our model—essentially a hierarchical Bayesian extension of phylogenetic generalized linear models (Paradis, 2012)—does make a few assumptions with respect to the phylogenetic information used. First, we incorporate the phylogeny at the family level for Lepidoptera. This implicitly assumes equal branch lengths for species within families, which is clearly not the case. However, we were unable to find a fully resolved species-level phylogeny for the Lepidoptera with adequate coverage for many of the families in our dataset, and we did not want to exclude species that were not represented in the phylogeny. Second, by treating the phylogenetic correlation matrix as a fixed covariate, we are unable to fully account for uncertainty in the phylogenetic relationships among taxa. This is an important assumption that could affect our estimates and conclusions, but one that is often made in these kinds of studies (O’Meara, 2012; Paradis, 2012).

Despite these assumptions and limitations, we are confident in our results for two reasons. First, we used correlation matrices from phylogenies assuming both Brownian motion and OU processes, as well as a matrix assuming no phylogenetic correlation at all. Model selection via WAIC indicated that all of these models were in the same family of “best models” (Table 2), and their parameter estimates (and the estimates from model averaging) were all quite similar, suggesting phylogeny is not playing a large role in the prevalence of *Wolbachia* infection among lineages. Second—and more gratifying—is that the posterior predictive checks for all models were quite similar (Figure 2). More important, data simulated from our model parameters closely matched the empirical data, indicating our approach is doing an excellent job at modeling the underlying biological process.

There are a number of interesting implications of our results. If the infection frequency for species is on the order of 12%, then perhaps *Wolbachia* is not presently a major player in the reproductive manipulation of this order. It follows that—if *Wolbachia* infection frequency is relatively low in the Lepidoptera—its role in the evolution of the order may not be as significant as with other groups (Miller et al., 2010). Accumulating evidence is demonstrating that *Wolbachia* is not an obligate manipulator of a host’s reproductive biology (Hamm et al., 2014a; Zhang et al., 2010, 2013). Indeed, Prout (1994) demonstrated that reproductive manipulator microbes should evolve to minimize harm to its host. Perhaps the paradigm needs to be reevaluated.

The assumption that *Wolbachia* always acts as a reproductive manipulator is clearly incorrect (Nice et al., 2009; Hamm et al., 2014b), and one should take care when extrapolating from the results of a positive *Wolbachia* assay to the real world. Luckily, this assumption is fading away. A *Wolbachia* infection can impart benefits to its host—for example the *wSuz* infection of *Drosophila suzukii* confers resistance to certain viruses (Cattel et al., 2016), can provide nutrition to its host (Hosokawa et al., 2010), and does not induce a manipulative phenotype (Hamm et al., 2014a). Thus, *Wolbachia* detection does not and should not imply detrimental effects to the host. Furthermore, our knowledge of *Wolbachia* as a reproductive manipulator in the Lepidoptera is based on scant evidence. To the best of our knowledge, of the 163 species of Lepidoptera considered positive for *Wolbachia*, only seven species from four families have been assayed for an induced phenotype (Table 1). Reciprocal cross experiments are required to determine what—if any—effects *Wolbachia* have on host reproduction. Until these experiments are conducted for a particular system we urge extreme caution when interpreting a positive PCR assay, and we hope that researchers will conduct the necessary experiments to determine if a manipulative phenotype exists within a particular system.

In many respects, the science of microbes in the insects, especially with regards to the endosymbiont *Wolbachia* and the Lepidoptera, is still in its natural history phase wherein research is largely in the descriptive phase, and as such we urge caution when interpreting positive *Wolbachia* assays and extrapolating consequences. Our model provides a framework to further our understanding of *Wolbachia* infection frequency in the Lepidoptera and, as we acquire more data, can generate new estimates. Ultimately, we hope to extend this model to the Insecta once sufficient data are acquired. We also consider that this model can be applied to other problems where one seeks to estimate infection frequency using hierarchical data.

## AUTHOR CONTRIBUTIONS

ZM and CH conceived of the experiment; ZM and CH conducted the analyses; ZM and CH wrote the manuscript.

## ACKNOWLEDGMENTS

We wish to thank James Fordyce, Ben Fitzpatrick, Christopher Peterson, Brian O’Meara and Jeremy Beaulieu for their input and advice during the preparation of this manuscript. We also wish to thank Francis Jiggins and Jack Welch for fruitful discussions while this project was nascent.

## SUPPLEMENTAL DATA

Supplementary Material should be uploaded separately on submission, if there are Supplementary Figures, please include the caption in the same file as the figure. LaTeX Supplementary Material templates can be found in the Frontiers LaTeX folder

